# Population genomic analysis uncovers African and European admixture in *Drosophila melanogaster* populations from the southeastern United States and Caribbean Islands

**DOI:** 10.1101/009092

**Authors:** Joyce Y. Kao, Asif Zubair, Matthew P. Salomon, Sergey V. Nuzhdin, Daniel Campo

## Abstract

Genome sequences from North American *Drosophila* melanogaster populations have become available to the scientific community. Deciphering the underlying population structure of these resources is crucial to make the most of these population genomic resources. Accepted models of North American colonization generally purport that several hundred years ago, flies from Africa and Europe were transported to the east coast United States and the Caribbean Islands respectively and thus current east coast US and Caribbean populations are an admixture of African and European ancestry. Theses models have been constructed based on phenotypes and limited genetic data. In our study, we have sequenced individual whole genomes of flies from populations in the southeast US and Caribbean Islands and examined these populations in conjunction with population sequences from Winters, CA, (USA); Raleigh, NC (USA); Cameroon (Africa); and Montpellier (France) to uncover the underlying population structure of North American populations. We find that west coast US populations are most like European populations likely reflecting a rapid westward expansion upon first settlements into North America. We also find genomic evidence of African and European admixture in east coast US and Caribbean populations, with a clinal pattern of decreasing proportions of African ancestry with higher latitude further supporting the proposed demographic model of Caribbean flies being established by African ancestors. Our genomic analysis of Caribbean flies is the first study that exposes the source of previously reported novel African alleles found in east coast US populations.

## Introduction

Out of the thousands of species in the genus *Drosophila*, the single most extensively studied species is *Drosophila melanogaster* (Powell 1997). The utility of *D. melanogaster* as a model organism can be seen in many fields of research from medicine to evolutionary biology. To fully take advantage of *D. melanogaster* as a model, we need the precision estimates and the history of population admixture during the species colonization of North America. The advent of next-generation sequencing (NGS), enabling the high-throughput sequencing of genomes, has generated much interest in the population genomics of *D. melanogaster* (Mackay *et al*. 2012; Pool *et al*. 2012; Campo *et al*. 2013) because understanding the population structure of *D. melanogaster* can now be approached with whole genome data (Duchen *et al*. 2013).

According to the currently accepted demographic model, *D. melanogaster* originated in sub-Saharan Africa with a migration event into the European continent 10,000 years ago (David & Capy 1988). Colonization of the Americas is hypothesized to have happened in two waves. The first wave occurred ∼400-500 year ago with African flies being transported into the Caribbean Islands along with the transatlantic slave trade. The second wave, which happened in the mid-19th century, was the cosmopolitan flies arriving with the first European settlers into North America (David & Capy 1988). These two waves purportedly created a secondary contact zone in the southeast United States and Caribbean Islands of cosmopolitan-adapted flies from Europe and African-like flies from West Africa (Caracristi & Schlötterer 2003; Duchen *et al*. 2013). The flies originating from the Caribbean islands have retained African-like behavior and physical phenotypes despite its close proximity to the US cosmopolitan populations (Yukilevich & True 2008a; Yukilevich & True 2008b; Yukilevich *et al*. 2010).

Previous studies looking at genome-wide effects of divergence in these populations used tiling microarrays to detect highly differentiated regions between the pooled genomes of cosmopolitan populations (including Caribbean fly lines) and Zimbabwean populations and then sequenced a subset of fragments to look at genetic divergence (Yukilevich *et al*. 2010). Most differentiation was found between populations living in African versus out of Africa and evidence supporting that most of the variation in North America and African populations originated from the sorting of African standing genetic variation into the New World through Europe (Yukilevich *et al*. 2010). However, Caracristi and Schlötterer (2003) found high levels of polymorphisms in North American populations where the proportion of shared alleles between African and American populations were greater than the proportion of shared alleles between African and European populations. This evidence supports the hypothesis that there was a separate migration event to the Caribbean and that this might be the source of these putative African alleles in North America (Li & Stephan 2006). More recently, Duchen et al. (2013) showed that North American populations of *D. melanogaster* are most likely the result of an admixture event between European and African populations with the African ancestry accounting for 15% of the mixture. However, it is not clear from their study whether there was a second migration event to the Caribbean from Africa. The Caribbean islands have been claimed to be the source of additional African alleles in the North American populations (Caracristi & Schlötterer 2003) although it has never been confirmed.

For this work, we have sequenced 23 *D. melanogaster* genomes from various locations in the southeast United States and the Caribbean Islands. Combined with the current sequencing efforts of other fly populations from Raleigh (NC, USA), Winters (CA, USA), Montpellier (France), and Oku (Cameroon), we can explore African and European admixture of North American populations in an attempt to elucidate the history of *D. melanogaster*’s migration to the Americas and to understand how Caribbean *D. melanogaster* populations can retain African-like phenotypes while being in such close proximity to European-like neighboring populations from the United States.

## Materials and Methods

### Fly Lines for Sequencing

A subset of 23 isofemale lines of *D. melanogaster* from 12 locations used in Yukilevich and True 2008b were selected for sequencing. Origins are as following: Selba, AL (ID#: 20, 28 and 20, 17); Thomasville, GA (ID#: 13, 34 and 13, 29); Tampa Bay, FL (ID#: 4, 12 and 4, 27); Birmingham, AL (ID#: 21, 39 and 21, 36); Meridian, MS (ID#: 24, 2 and 24, 9); Sebastian, FL (ID#: 28, 8); Freeport, Grand Bahamas-west (ID#: 33, 16 and 33, 11); George Town, Exumas (ID#: 36, 9 and 36, 12); Bullock’s Harbor, Berry Islands (ID#: 40, 23 and 40, 10); Cockburn Town, San Salvador (ID#: 42, 23 and 42, 20); Mayaguana, Mayaguana (ID#: 43, 19 and 43, 18); Port Au Prince, Haiti (ID#: H, 29 and H, 25). All flies were maintained at 25 °C in vials on a standard cornmeal diet.

### Libraries and sequencing of southeast US and Caribbean lines

All lines were subjected to full-sibling inbreeding for at least five generations before we collected 15 - 20 females from each line for library preparation. DNA was extracted using a Epicentre MasterPure kit (Madison, WI, USA) and cleaned with the Zymo Quick-gDNA Miniprep kit (Irvine, CA, USA). Illumina sequencing libraries were prepared according to Dunham and Friesen (2013) with the exception that DNA was sheared with dsDNA Shearase Plus (Zymo: Irving, CA, USA) and cleaned using Agencourt AMPure XP beads (Beckman-Coulter: Indianapolis, IN, USA). Fragment size selection was also done using beads instead of gel electrophoresis. Libraries were visualized in an Agilent Bioanalyzer 2100 and quantified using the Kapa Biosystems Library Quantification Kit, according to manufacturer’s instructions. Libraries were loaded into an Illumina flow cell v.3 and run on a HiSeq 2000 for 2×100 cycles. Library quality control and initial sequencing were performed at the USC NCCC Epigenome Center’s Data Production Facility (University of Southern California, Los Angeles, CA, USA). Additional sequencing to achieve at least 5x genome-wide coverage for all lines was performed at the USC UPC Genome and Cytometry Core (University of Southern California, Los Angeles, CA, USA), in an Illumina HiSeq 2500 following the same run format.

### Sources of other sequenced populations

We used the 35 isogenic lines from Winters, CA, USA and 33 isogenic lines from Raleigh, NC, USA described in Campo *et al*. (2013). Raleigh lines were a subset of the Drosophila Genetic Reference Panel (DGRP) (Mackay et al, 2012). The 10 isofemale lines from Oku, Cameroon, were sequenced as a part of the Drosophila Population Genetic Panel (DPGP-2 African Survey) (Pool *et al*. 2012). Sequencing reads for 20 isofemale lines from Montpellier, France were downloaded via the Bergman lab webpage (Haddrill & Bergman 2012).

### Mapping

For each fly line, the raw sequencing reads were trimmed by quality using the SolexaQA package (ver. 1.12) with default parameters and all trimmed reads less than 25 bp were discarded (Cox *et al*. 2010). The quality trimmed reads were then mapped to the *D. melanogaster* reference genome (FlyBase version 5.41) using Bowtie 2 (ver. beta 4) with the “very sensitive” and “-N=1” parameters (Salzberg & Langmead 2012). Following mapping, the GATK (ver. 1.1-23, dePristo *et al*. 2011) IndelRealigner tool was used to perform local realignments around indels and PCR and optical duplicates were identified with the MarkDuplicates tool in the Picard package (http://picard.sourceforge.net).

### SNP calling, phasing, and filtering

SNP variants were identified in all lines simultaneously using the GATK UnifiedGenotyper (ver. 2.1-8) tool with all parameters set to recommended default values. The raw SNP calls were further filtered following the GATK best practices recommendations (Auwera *et al*. 2013) resulting in 4,021,717 SNP calls. We then used BEAGLE to perform haplotype phasing as well as impute missing data (Browning & Browning 2007; Browning & Browning 2009). SNPs were further filtered using VCFtools (http://vcftools.sourceforge.net/) for 5% minor allele frequency and biallelic sites resulting in 1,047,913 SNPs across the major chromosomal regions: 2L (222,464 SNPs), 2R (192,120 SNPs), 3L (212,601 SNPs), 3R (268,701 SNPs), and X (152,027 SNPs) to be considered for further analysis.

### Population structure analysis

We used VCFtools (Danecek *et al*. 2011) to calculate F_ST_ via the Weir and Cockerham estimates (1984) as a proxy for genetic distance between all our populations. Additionally, we used the R package SNPRelate (Zheng *et al*. 2012) to perform principal component analysis (PCA). We did PCA with all populations and then removed the Cameroon population for another PCA to investigate North American patterns further without the influence of the African population.

ADMIXTURE (Alexander *et al*. 2009) estimates ancestry of a given set of unrelated individuals in a model-based manner from large autosomal SNP genotype datasets. The program outputs the proportion of ancestral population for each individual. To run the program, a prior belief number of ancestral populations (K), must be provided. We used a cross-validation procedure of ADMIXTURE to propose the number of ancestral populations (K). Optimal K values will have lower cross-validation error compared to other values. We ran a 5-fold cross validation on the plink file (.ped) which was generated using a custom PERL script from the Variant Calling File (VCF). Linkage disequilibrium can affect the results of ADMIXTURE thus the marker set used for this analysis was further filtered to include only autosomal markers that were at least 250 bp apart resulting in a total of 234,497 SNPs.

### Chromosome painting

We utilized the software Chromopainter (Lawson *et al*. 2012) to estimate which parts of the genome each North American individual were contributed by European or African ancestors. We ran Chromopainter for 60 iterations to estimate parameters of the algorithm and then ran Chromopainter with the estimated parameters to obtain the final results as recommended in the user manual. Additionally, we implemented hierarchical clustering in R (heatmap.2 with standard options in the gplots library) to examine the similarity of Chromopainter results across each chromosomal region between all the North American individuals.

### Linkage Disequilibrium Analysis

To look at linkage disequilibrium decay over genomic distance, measures of D’ were estimated using VCFtools (Danecek *et al*. 2011) in 10,000 bp windows across the genome.

## Results

### Investigating Population Structure by Principal Component Analysis

To explore initial relationships between populations, we performed PCA on the 1,047,913 quality-filtered SNPs using the R package SNPRelate. The first principal component represented the separation between African and non-African populations and the second principal component was the variation within the Cameroon population (FIGURE 2). Upon closer inspection of the non-African cluster (FIGURE 2), the first principal component could also be a proxy to how genetically close each non-African population is to the Cameroon population, with the Caribbean population located the closest. The non-African populations were roughly grouped into two sub-clusters of Caribbean and non-Caribbean. There were, however, a few Caribbean fly lines that clustered close to and within the non-Caribbean group. The four Caribbean lines that clustered with the US populations were collected from locations on islands closest to the US and Caribbean border (i.e. Freeport, Grand Bahamas-west and Bullock’s Harbor, Berry Islands). Along with these four Caribbean lines, the sequenced fly lines from locations in the southeast United States were interspersed with fly lines from Raleigh, indicating a potential east coast US admixture zone. The Raleigh population clustered very closely with the Winters, but both Raleigh and Winters appeared to still be distinct populations. The 20 French lines appeared dispersed in the non-Caribbean cluster, which supports the notion that there is much European influence in North American populations.

**FIGURE 1.**
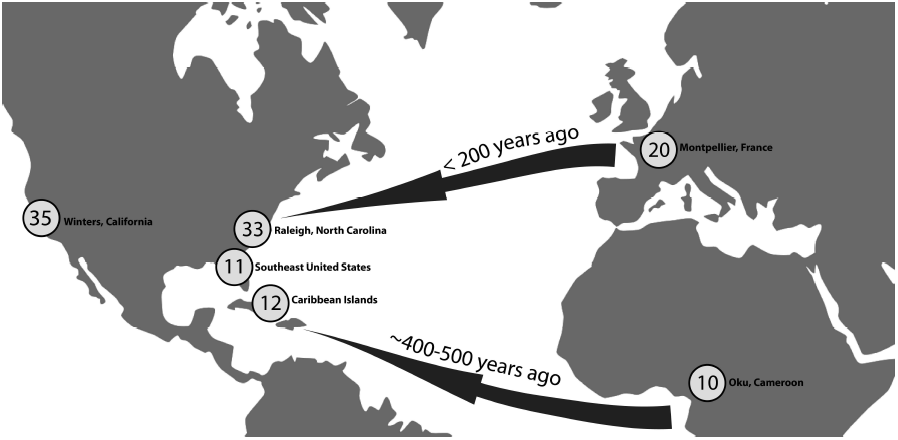
Map of sequenced populations with number of whole genome sequences in circles. Arrows indicate currently accepted migration history of *D. melanogaster* into the Americas.

**FIGURE 2.**
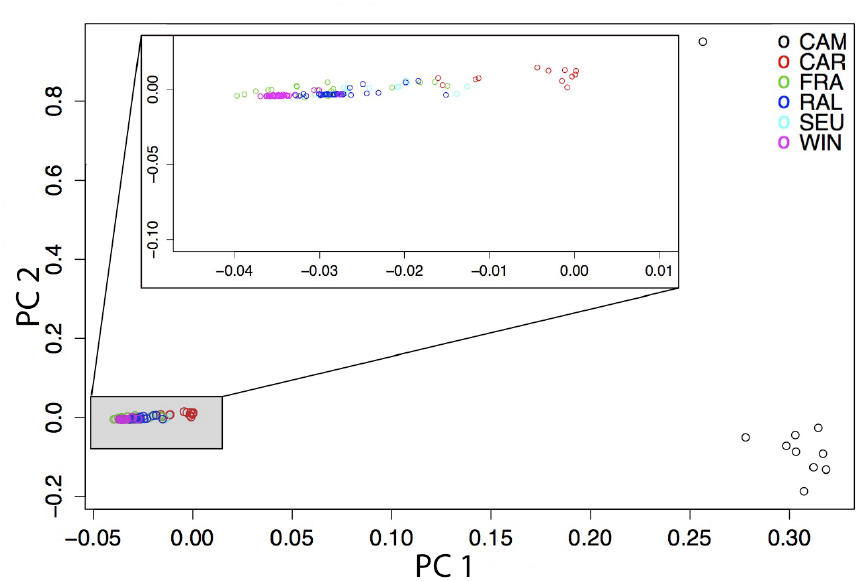
First and second principal components (PC) from principal components analysis with populations from Cameroon (CAM), Caribbean Islands (CAR), France (FRA), Raleigh (RAL), southeast US (SEU) and Winters (WIN). Population structure of individuals in the grey highlighted box are magnified in secondary enlarged plot.

Upon inspection of additional principal components (FIGURE S1), principal components 3 and 4 explained variation within the Cameroon population indicating there was much diversity in the African population, which may have been masking patterns in the non-African populations. We removed the Cameroon population and performed a second PCA using non-African populations (FIGURE S2). The first principal component in this second PCA explained the variation within the North American populations, while the second principal component separated the French population from the North American populations. Clustering patterns of the second PCA were similar to those in the first PCA, but we saw that the French population formed a distinct cluster and was located closest to the group containing Winters, Raleigh, and southeast US populations. The third and fourth principal components accounted for more variation within the North American populations (FIGURE S3).

### Genetic differentiation between populations

To quantify the level of genetic differentiation, we calculated Weir and Cockerham (1984) F_ST_ between all pairs of populations per SNP and averaged the F_ST_ estimates per chromosomal region. We found a consistent pattern in which Cameroon was highly differentiated from all cosmopolitan populations, but was closest to the Caribbean population (FIGURE 3). The French and Winters populations were the most differentiated from the Cameroon lines. As expected, the greatest differentiation between the Cameroon population and the non-African populations was on the X chromosome (FIGURE 3), since this chromosome has been suggested to evolve faster than the autosomes (Presgraves 2008).

**FIGURE 3.**
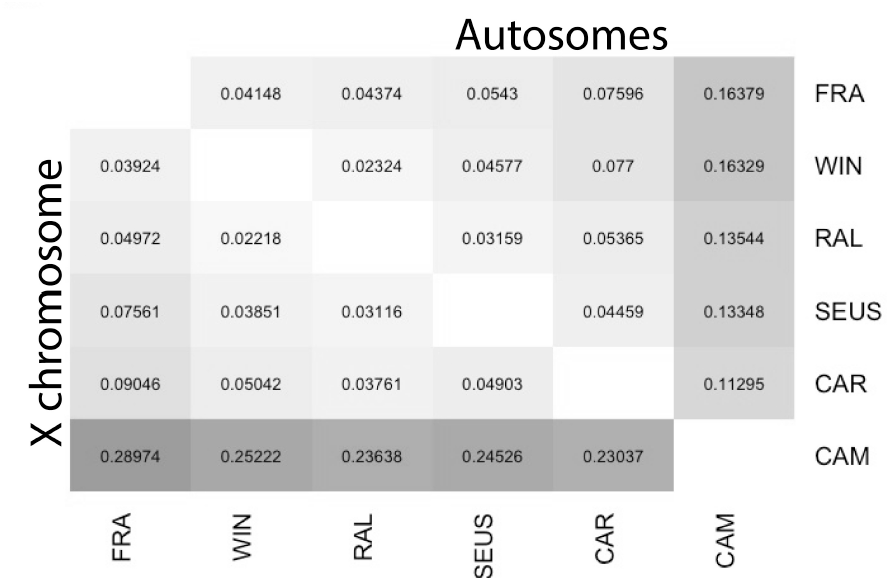
Average F_ST_ values between populations for chromosome X (lower diagonal) and all autosomes (upper diagonal). Shades of grey illustrate the degree of genetic differentiation with larger F_ST_ values being darker and smaller F_ST_ values being lighter.

The French population was the least genetically differentiated from the Winters and Raleigh populations (FIGURE 3). Interestingly enough, the Caribbean population was slightly more differentiated from the Winters population than from the French population in the 2L and 3R chromosomal regions (Supplementary TABLE 1,2), perhaps indicating a slightly larger European influence in the Caribbean than the west coast US.

### Admixture patterns

From our cross-validation procedure, it was determined that the optimal number of ancestral populations for ADMIXTURE was K=2 (FIGURE S4). According to the ancestral proportions (FIGURE 4A), it appears that the North American lines are a composite of European and African ancestry. Furthermore, the proportion of African-like markers is higher in Caribbean individuals and decrease in proportion with increasing latitude (FIGURE 4B).

**FIGURE 4.**
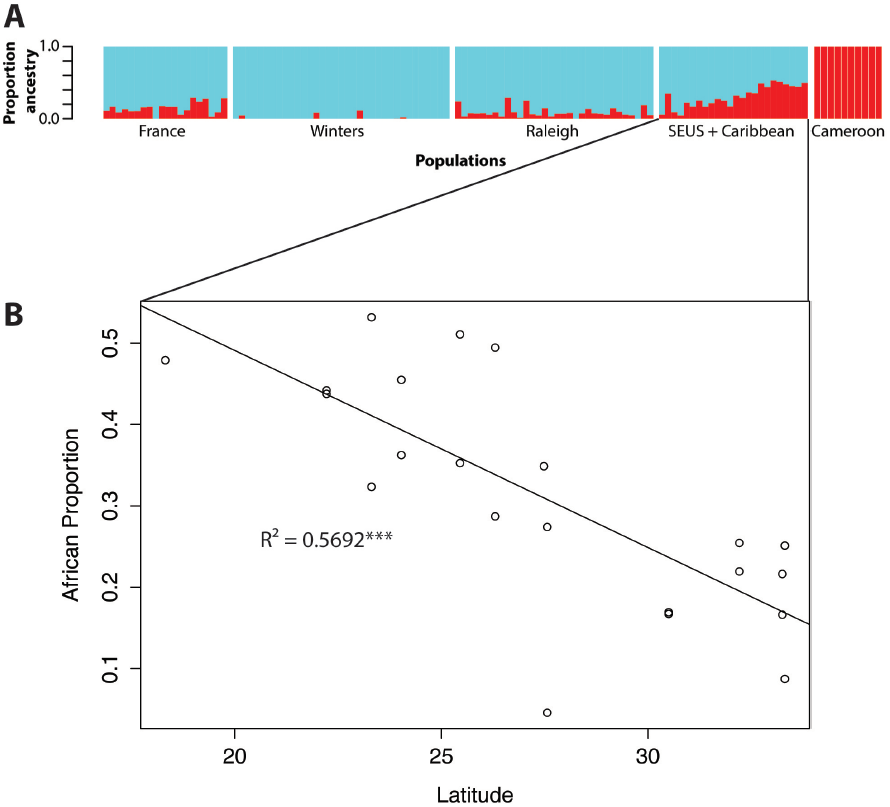
A) ADMIXTURE results of quality and LD filtered autosomal markers for two ancestral populations (K=2). B) Relationship between latitude and proportion of African ancestry of southeast US and Caribbean individuals. Asterisks on the R^2^=0.5692 corresponds with F=26.42 and a significance of P < 0.0001.

### Genome-wide African and European influences

While results from ADMIXTURE are useful in understanding how populations are structured and point towards approximate the influences of African and European ancestors, we cannot determine the pattern of influence across a genome with those results. We used Chromopainter to predict the ancestry of all the North American sequenced fly lines across the genome. The most striking result from visualizing the local ancestry of all genomes (FIGURE 5) was that larger chunks of African or European ancestry seemed to be retained in telomeric and centromeric regions known to have low recombination (Comeron *et al*. 2012).

**FIGURE 5.**
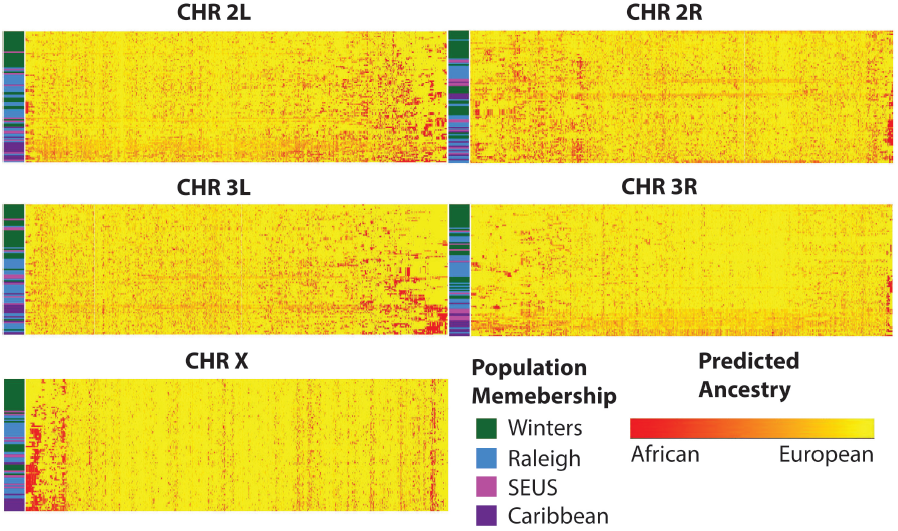
Painted chromosomal regions heatmap with hierarchical clustering of individuals. Each row in heatmap represents one individual. Population membership of individual designated by vertical bar to the right of each chromosomal heatmap (Green: Winters, CA, Blue: Raleigh, NC, Pink: Southeast US, Purple: Caribbean). Red represents SNPs that are most similar to the Cameroon donor population; Yellow represents SNPs that are most similar to the French donor population.

When we clustered individual genomes by genomic inheritance patterns, the patterns of individuals within one population clustered more with each other than with other populations except for chromosomal region 2R where Caribbean and southeast US individuals seem to be evenly dispersed between Winters and Raleigh populations. Chromosome X appeared to be the least influenced by African ancestry (FIGURE 5), which is in agreement with the large X effect (Presgraves 2008).

Individuals from the Caribbean populations and some from the southeast US seemed to have a larger percentage of African painted alleles, which was especially apparent in the chromosomal regions of 2L and 3R (FIGURE 5). The long stretches of the African-painted SNPs in these chromosomal regions coincided with the locations of common cosmopolitan inversions, In(2L)t and In(3R)P (Corbett-Detig & Hartl 2012).

Overall the expected proportion of probable African ancestry ranged between 3.6% (Winters, CA) to 47% (Caribbean Islands) for the painted genomes. On average over the whole genome, the expected percentage of African ancestry was highest in the Caribbean population at 24.75% and the lowest in the Winters population at 8.68%. Raleigh and southeast US populations had 14% and 15.6% of predicted African ancestry, which is consistent with previous findings (Duchen *et al*. 2013). In summary, populations had decreasing African ancestry with respect to distance from the Caribbean Islands in all genomic areas. Out of all the chromosomes, the X chromosome had the lowest expected percentage of African-inherited alleles for all North American populations (FIGURE S5).

### Linkage disequilibrium patterns

Elevated levels of linkage disequilibrium (LD) can be an indicator of admixture in populations because inherited ancestral tracts have not had sufficient time to be broken down by recombination (Loh *et al*. 2013). We calculated D’ as a measure of LD and averaged the absolute value of D’ to get approximate LD levels in our populations across different genomic regions. We found that on average Cameroon and France populations have lower LD values than North American populations (FIGURE 6). Out of all the North American populations, the Caribbean population had one of the lowest LD values on most chromosomal regions except on the X chromosome. This is consistent with the notion that African flies colonized the Caribbean Islands a good 200 years before European flies arrived on the east coast of the US making the Caribbean population older than the US populations (David & Capy 1988).

**FIGURE 6.**
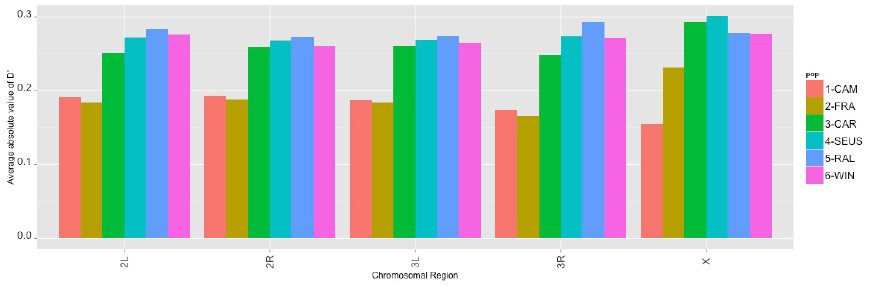
Average |*D*′| as a measure of linkage disequilibrium by population and chromosome

## 4.4 Discussion

### Caribbean flies most likely established by African ancestors

Although all non-African populations pairwise F_ST_ values were high throughout the genome when compared to the African sample, the Caribbean population had on average the lowest values. With the Caribbean population located closest in the first PC analysis to the Cameroon population and the highest percentage of predicted African ancestry out of all the North American samples we analyzed, these pieces of evidence do seem to further support the migration event of west African flies to the Caribbean islands via the transatlantic slave trade (David & Capy 1988).

### African and European admixture in North America

Recently admixed populations exhibit more linkage disequilibrium than older long-established populations (Loh *et al*. 2013). This is because newer populations, which are a combination of genetic material from older base populations have not gone through enough generations for recombination to break down LD blocks. We do detect higher LD in the North American populations than in our African and European samples. Although this is a common signature of admixture, higher LD values can also result from other demographic events such as a population bottleneck. However, previous studies have already established the existence of admixture in some North American populations, particularly Raleigh, (Duchen *et al*. 2013) which would support that elevated LD in our case is most likely due to admixture.

We are able to extend the admixture scenario in North America with our 23 sequenced genomes from the southeast US and Caribbean islands. It has been postulated that American *D. melanogaster* are more genetically variable than European *D. melanogaster* due to admixture from the Caribbean islands (Caracristi & Schlötterer 2003). Our results from ADMIXTURE (FIGURE 4) and chromosome painting (FIGURE 5) clearly show a clinal pattern of African introgression into the United States, which supports the notion that these non-European African alleles in the US are originating from the Caribbean Islands. Furthermore, the PCA groupings (FIGURE 2) also illustrate that the border between the southeast US and Caribbean Islands is where fly populations are experiencing the most admixture.

### Westward expansion of Drosophila melanogaster

Our analysis of the Winters, CA genomes revealed that the Winters population is more related to our European population than the other US population. There appears to be very little to no African ancestry in the genomes from Winters, CA. Either there was a separate colonization event in the west or when *D. melanogaster* arrived in North America with European settlers, it quickly expanded west shortly after arriving (Campo *et al*. 2013). The latter explanation may be more plausible given that the first sighting of *D. melanogaster* was in the mid-19th century (David & Capy 1988), which was when the United States was in the midst of active westward expansion with the rapid construction of a transcontinental railway to transport supplies out to early settlers in the west (Billington 1949).

## Conclusions

Understanding the origins and genomic patterns of North American *D. melanogaster* will be useful for researchers working with populations from this area of the world especially with the emerging public sequencing data becoming available (Mackay *et al*. 2012; Remolina *et al*. 2012). Our genome analyses of southeast US and Caribbean fly populations in relation to other North American populations and to their African and European ancestral populations further elucidate the history of *Drosophila melanogaster* colonization of North America. We reveal clinal patterns of African ancestry from the Caribbean Islands to the southeast United States illustrating African and European admixture maintained in those populations, which is likely influencing populations that lie farther north on the east coast of the United States.

## Chapter Bibliography

Alexander DH, Novembre J, Lange K (2009) Fast model-based estimation of ancestry in unrelated individuals. Genome Research, 19, 1655–1664.

Auwera GA, Carneiro MO, Hartl C et al. (2013) From FastQ Data to High-Confidence Variant Calls: The Genome Analysis Toolkit Best Practices Pipeline. John Wiley & Sons, Inc., Hoboken, NJ, USA.

Billington RA (2001) The Transportation Frontier. In: Westward expansion: a history of the American Frontier 6^th^ ed., abridged, pp. 279–198. University of New Mexico Press, Albuquerque, NM.

Browning BL, Browning SR (2009) A Unified Approach to Genotype Imputation and Haplotype-Phase Inference for Large Data Sets of Trios and Unrelated Individuals. The American Journal of Human Genetics, 84, 210–223.

Browning SR, Browning BL (2007) Rapid and Accurate Haplotype Phasing and Missing-Data Inference for Whole-Genome Association Studies By Use of Localized Haplotype Clustering. The American Journal of Human Genetics, 81, 1084–1097.

Campo D, Lehmann K, Fjeldsted C et al. (2013) Whole-genome sequencing of two North American Drosophila melanogasterpopulations reveals genetic differentiation and positive selection. Molecular Ecology, 22, 5084–5097.

Caracristi G, Schlötterer C (2003) Genetic differentiation between American and European Drosophila melanogaster populations could be attributed to admixture of African alleles. Molecular Biology and Evolution, 20, 792.

Cockerham C (1984) Estimating F-statistics for the analysis of population structure. Evolution.

Comeron JM, Ratnappan R, Bailin S (2012) The Many Landscapes of Recombination in Drosophila melanogaster. PLoS Genetics, 8, e1002905.

Corbett-Detig RB, Hartl DL (2012) Population Genomics of Inversion Polymorphisms in Drosophila melanogaster. PLoS Genetics, 8, e1003056.

Cox MP, Peterson DA, Biggs PJ (2010) SolexaQA: At-a-glance quality assessment of Illumina second-generation sequencing data. BMC bioinformatics, 11, 485.

Danecek P, Auton A, Abecasis G et al. (2011) The variant call format and VCFtools. Bioinformatics, 27, 2156–2158.

David J, Capy P (1988) Genetic variation of Drosophila melanogaster natural populations. Trends in genetics : TIG, 4, 106–111.

DePristo MA, Banks E, Poplin R et al. (2011) A framework for variation discovery and genotyping using next-generation DNA sequencing data. Nature Genetics, 43, 491–498.

Duchen P, Živković D, Hutter S, Stephan W, Laurent S (2013) Demographic Inference Reveals African and European Admixture in the North American Drosophila melanogaster Population. Genetics, 193, 291–301.

Dunham JP, Friesen ML (2013) A Cost-Effective Method for High-Throughput Construction of Illumina Sequencing Libraries. Cold Spring Harbor Protocols, 2013, pdb.prot074187–pdb.prot074187.

Jeffrey R. Powell Department of Biological Sciences Yale University (1997) Progress and Prospects in Evolutionary Biology : The Drosophila Model. Oxford University Press.

Langmead B, Salzberg SL (2012) Fast gapped-read alignment with Bowtie 2. Nature methods, 9, 357–359.

Lawson DJ, Hellenthal G, Myers S, Falush D (2012) Inference of Population Structure using Dense Haplotype Data. PLoS Genetics, 8, e1002453.

Li H, Stephan W (2006) Inferring the demographic history and rate of adaptive substitution in Drosophila. PLoS Genetics, 2, e166.

Loh P-R, Lipson M, Patterson N et al. (2013) Inferring Admixture Histories of Human Populations Using Linkage Disequilibrium. Genetics, 193, 1233–1254.

Mackay TFC, Richards S, Stone EA et al. (2012) The Drosophila melanogaster Genetic Reference Panel. Nature, 482, 173–178.

Pool JE, Corbett-Detig RB, Sugino RP et al. (2012) Population Genomics of Sub-Saharan Drosophila melanogaster: African Diversity and Non-African Admixture. PLoS Genetics, 8, e1003080.

Presgraves DC (2008) Sex chromosomes and speciation in Drosophila. Trends in Genetics, 24, 336–343.

Pritchard J, Stephens M, Donnelly P (2000) Inference of population structure using multilocus genotype data. Genetics, 155, 945.

Remolina SC, Chang PL, Leips J, Nuzhdin SV, Hughes KA (2012) Genomic basis of aging and life-history evolution in Drosophila melanogaster. Evolution, 66, 3390–3403.

Yukilevich R, True JR (2008a) INCIPIENT SEXUAL ISOLATION AMONG COSMOPOLITAN DROSOPHILA MELANOGASTER POPULATIONS. Evolution, 62, 2112–2121.

Yukilevich R, True JR (2008b) AFRICAN MORPHOLOGY, BEHAVIOR AND PHERMONES UNDERLIE INCIPIENT SEXUAL ISOLATION BETWEEN US AND CARIBBEAN DROSOPHILA MELANOGASTER. Evolution, 62, 2807–2828.

Yukilevich R, Turner TL, Aoki F, Nuzhdin SV, True JR (2010) Patterns and Processes of Genome-Wide Divergence Between North American and African Drosophila melanogaster. Genetics, 186, 219–239.

Zheng X, Levine D, Shen J et al. (2012) A high-performance computing toolset for relatedness and principal component analysis of SNP data. Bioinformatics, 28, 3326–3328.

